# Teach your microscope how to print: Low-cost and rapid-iteration microfabrication for biology

**DOI:** 10.1101/2025.02.20.639256

**Authors:** Lucien Hinderling, Remo Hadorn, Moritz Kwasny, Joël Frei, Benjamin Grädel, Sacha Psalmon, Yannick Blum, Rémi Berthoz, Alex Landolt, Benjamin D. Towbin, Daniel Riveline, Olivier Pertz

## Abstract

The application of traditional microfabrication techniques to biological research is hindered by their reliance on clean rooms, expensive or toxic materials, and slow iteration cycles. We present an accessible microfabrication workflow that addresses these challenges by integrating consumer 3D printing techniques and repurposing standard fluorescence microscopes equipped with DMDs for maskless photolithography. Our method achieves micrometer-scale precision across centimeter-sized areas without clean room infrastructure, using affordable and readily available consumables. We demonstrate the versatility of this approach through four biological applications: inducing cytoskeletal protrusions via 1 μm-resolution surface topographies; micropatterning to standardize cell and tissue morphology; fabricating multilayer microfluidic devices for confined cell migration studies; imprinting agar chambers for long-time tracking of *C. elegans*. Our protocol drastically reduces material costs compared to conventional methods and enables design-to-device turnaround within a day. By leveraging open-source microscope control software and existing lab equipment, our workflow lowers the entry barrier to micro-fabrication, enabling labs to prototype custom solutions for diverse experimental needs while maintaining compatibility with soft lithography and downstream biological assays.

## Introduction

Microfabrication technologies using photolithography and soft lithography have enabled researchers to build cellular environments with micrometer precision. By shaping and patterning the geometry, topology and composition of the extracellular space at a precision that matches the intrinsic scale of cells, these technologies provide a powerful tool to study interactions between cellular systems and their environment [1–3]. Microfabricated structures can be used in a wide array of downstream applications: to confine bacteria [4], eukaryotic cells [5–7], or microscopic animals [8]; fabricate microfluidic devices [9] e.g. to study 3D migration [10, 11], or mimic complex geometries to grow more realistic organ [12] and cancer [13, 14] models in vitro; to pattern surfaces to study 2D cell migration [15–19], measure forces exerted by cells onto their substrate [20], and homogenize cell morphologies [21–24]. Furthermore, lab-on-a-chip devices use microfabrication to miniaturize systems, enabling massive parallelization [25]. Photolithography techniques for biological research were originally adapted from the semiconductor industry [26, 27]. These methods were designed to meet the sub-micrometer precision requirements of electronics manufacturing, whereas many biological applications do not require such precise spatial resolution. Bottlenecks in microfabrication processes are that they require clean room access [28, 29] and expensive single-purpose hardware such as mask aligners. Further, the required reagents and substrates can be costly, toxic, and difficult to source. The continued reliance on high-precision techniques for biological research has led to unnecessary complexity, creating barriers to accessibility [30, 31]. Traditionally photolithography uses a high resolution photomask that contains the design. Although commercially available photomasks are relatively inexpensive, their long production times hinder iterative design cycles. Recent maskless approaches and efforts to simplify protocols are improving iteration speed and accessibility of these methods [18, 30, 32–35]. This approach has been commercialized by Alvéole is widely used.

In contrast, 3D printing has significantly expanded access to additive manufacturing and rapid prototyping, proving useful in many laboratories [36, 37]. A class of 3D printers recently becoming available for the consumer market, commonly known as “resin printers”, employs photolithographic techniques to achieve x/y resolutions down to 50 μm [31], but they still fall short of producing geometric features at the cellular scale.

By combining elements of semiconductor microfabrication with consumer 3D printing, we present a simplified microfabrication protocol tailored for biological applications, achieving micrometer precision at the centimeter scale while reducing time and procedural complexity. We drastically reduce the cost of consumables by replacing the commonly used SU-8 photoresist with 3D printing resin, and silicon wafers with standard microscope slides. Our approach repurposes an existing microscope setup used for targeted photostimulation as a maskless microfabrication system, streamlining the process from concept to fabricated structure within a day. Compared to commercial solutions, our approach relies on open-source software and does not require specific proprietary hardware or reagents, ensuring compatibility with existing equipment, reducing costs, and enabling customization.

We first describe the method, then demonstrate its application across various biological model systems and scales, ranging from subcellular to whole organisms, and from micrometers to centimeters. Specifically, we demonstrate:

1. Fabrication of μm-scale pillar topologies to guide formation of cytoskeletal structures.
2. Surface patterning with adhesive and non-adhesive coatings to control and standardize fibroblast cytoskeletal organization and 2D gastruloid growth.
3. Manufacturing of microfluidic devices to study confined cell migration through constrictions.
4. Imprinting chambers into agar to confine *C. elegans* movement.

## Results

### Simplified rapid iteration microfabrication workflow

We first create a microfabricated structure using maskless photolithography (fig. 1A). In this process, an image mask designed in a computer graphics software is projected onto a thin film of UV-curable resin. Like in a standard video projector, the light is shaped using a DMD, but instead of projecting the image onto a screen, the image is demagnified using a microscope objective and focused on the microscope slide. We can repurpose our microscope set up for targeted optogenetic photostimulation without any hardware modifications. In the areas exposed to UV light the resin hardens, while the unexposed regions remain soft and are washed away, leaving behind a mold that can be replicated with an elastomer such as polydimethylsiloxane (PDMS). The resulting PDMS copy serves as a foundation for downstream applications, such as microfluidics and stamping.

**Fig. 1.**
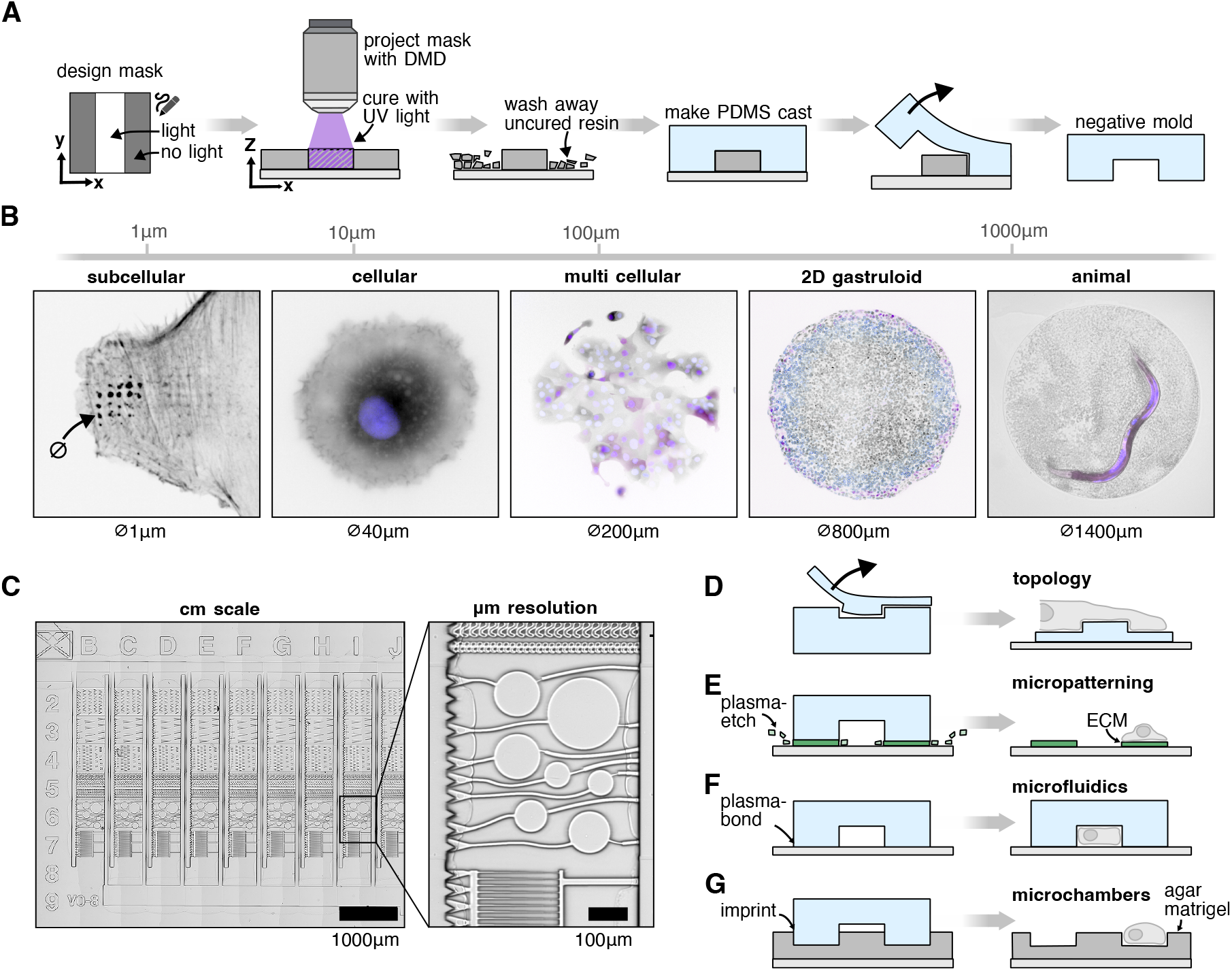
Microfabrication method. **A** Protocol overview. A mask is designed using computer graphics software. Using a microscope, UV light is projected onto a glass slide coated with UV curable resin. The uncured resin is then washed away and a PDMS cast is formed from the 3D printed structure. **B** Our method can be applied across biological scales, from subcellular to cellular. Subcellular: A REF52 cell forming protrusions into microfabricated wells (fig. 2, *lifeact::mNeonGreen*). Cellular: A single NIH3T3 cell on a circular fibronectin micropattern (fig. 3, black: *ERK-KTR::mRuby2*; blue: *H2B::miRFP670*)). Multi cellular: Sparsly seeded MCF10A cells on a circular fibronectin micropattern (black: *ERK-KTR::mRuby2*; blue: *H2B::miRFP670*). 2D gastruloid: Human embryonic E9 stem cells growing on a circular matrigel pattern (fig. S5, black: *H2B::miRFP670*; blue: *OCT4:POM121::tdTomato*; purple: Brachyury immunostain). Animal: *C. elegans* worm confined in a microwell (fig. 5, purple: *eft-3p::mScarlet*). **C** Structures at the cm scale can be fabricated while maintaining μm resolution (here a microfluidic chip, fig. 4). **D-F** The microfabrication technique can be used for a variety of applications: (D) patterning of surface topology with subcellular precision (fig. 2), (E) patterning of surface chemistry to e.g. induce a specific cell morphology (fig. 3), (F) microfluidic devices (fig. 4), (G) imprinting other substrates like matrigel or agar, using the PDMS mold as a stamp (fig. 5).

An overview of our method is provided here, with a detailed protocol and additional practical considerations outlined in the methods section of this paper. The required consumables and devices are provided in tables 1 and 2.

**Table 1.** Consumables microfabrication.

|  |  |  |  |
| --- | --- | --- | --- |
| **TMSPMA** 3-(Trimethoxysilyl)propylmethacrylate | Sigma Aldrich, 440159 | 100 ml | \$71 |
| **PDMS** Polydimethylsiloxane | SYLGARD 184 Silicone Elastomer Kit | 0.5 kg | \$162 |
| **UV-printing resin** | Copymaster3D Tough UV Resin Clear | 500 ml | \$31 |
| **Methanol** |  |  |  |
| **Isopropanol** |  |  |  |
| **Ethanol** |  |  |  |
| **Microscope slides** |  |  |  |

**Table 2.** Devices microfabrication.

|  |  |
| --- | --- |
| Microscope with DMD and μManager | Nikon TiE, Andor Mosaic 3 / Mightex Polygon 100 |
| UV light source | Lumencor Spectra X |
| Spin coater | Ossila 120 – 6,000 rpm |
| Plasma cleaner | Femto Science, Cute |
| Oven 70 °C |  |

The procedure begins by coating a standard microscope slide with 3-(Trimethoxysilyl)propylmethacrylat (TMSPMA) to enhance the adhesion of the UV resin [38]. A thin layer of consumer-grade 3D-printing resin is then spin-coated onto the slide. The prepared slide is placed onto a standard fluorescence microscope (Eclipse Ti, Nikon) equipped with a system capable of projecting UV light patterns. In our setup, we use a commercial DMD (Mosaic 3, Andor or Polygon 1000, Mightex) available as microscope attachment in combination with a 395 nm UV light source (Spectra X, Lumencor) to project patterns onto the resin. By adjusting the microscope objective, we can control the projected feature size: a 20× air objective offers a good tradeoff between field of view (FOV) size and resolution for most applications. For large features and thick layers, we use a 4x objective because of its increased depth of field. Fig. 1B illustrates applications of microfabricated circles ranging from 1 to 1400 μm. Using the microscope’s x/y stage, we iteratively project images to create patterns larger than the FOV while maintaining spatial resolution (fig. 1C). The UV exposure time is dependent on the layer height, objective and light source used, but can be empirically calibrated within 5 min (fig. S1). As the resin is very sensitive to UV light, we never exceeded illumination times of 1000 ms per FOV. A microscope slide can thus easily be scanned and exposed within a few minutes. The resin is designed for the consumer market, optimized for use in home settings without requiring specialized environments like yellow light rooms typically needed in conventional photolithography.

After exposure, the unexposed resin is washed away, forming the initial resin mold. This mold is then post-cured using UV light and heat [39]. Once cured, PDMS is poured over the mold and hardened in an oven. The PDMS cast is then lifted off and can be used for various experiments, including controlling surface topology, micropatterning surfaces, fabricating microfluidic devices, and creating agar microchambers (fig. 1D-F). We show results for these four applications in the next chapters. The theoretical maximal resolution of the projection for a certain optical setup can be calculated by measuring the FOV of the DMD divided by its resolution. For a 20× objective, we measure a projection width of 560 μm, divided by 800 px DMD x-resolution we get a resolution of ~0.7 μm/px. In practice, diffraction artifacts from the DMD mirror edges and the optical path are reducing the resolution. Using a 20× objective, we can reliably print features sizes down to 5 μm (fig. S2), smaller features are possible depending on the design. For the 1 μm sized pits in the first example, we used a 100× oil immersion objective. The footprint of the structure is limited by the carrier glass used (up to 50×70 mm minus border padding) or stage travel range. The z-layer height *h* is controlled by the spin coating rpm, and can be calculated with a calibration test print and the following formula:

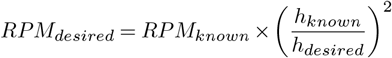

To calibrate for a large range of rpm (200-3200 rpm), we find that a slightly more complex model with a constant offset leads to a good fit of the data (R^2^: 0.96 with offset, R^2^: 0.90 without offset):

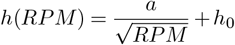

where *h* describes the film thickness as a function of the spin speed *RPM*, *a* is a proportionality constant and *h*_0_ represents a constant offset (supp. fig. S3).

To automate the printing process, we control the microscope using custom code available on GitHub^1^. We employ μManager [40] in combination with Pycromanager [41] to control the hardware from Python, and Napari [42] to visualize the camera feed and provide interactivity. An interactive notebook with step-by-step instructions simplifies the initial hardware setup and experiment execution, making it accessible even to researchers without coding experience.

### Engineering surface topology with 1 μm size features to guide cell protrusions

Cells can sense and respond to the geometry and topology of their 3D extracellular matrix surroundings. Early approaches to creating artificial topologies used natural materials such as spider webs [43] and fibrin fibers extracted from plasma [44]. More recently, microfabrication techniques have been employed to achieve greater control over surface structures [45–47].

Here, we create an artificial surface topology featuring round pits or pillars with diameters as small as 1 μm (fig. 2A). This is achieved by printing pillars using backside illumination with a 100× oil objective through a coverslip. The printed structure is cast into a negative PDMS stamp, forming a surface with pits. This negative is then cast again into a positive PDMS stamp to produce pillar structures. For experimental use, either the pit or pillar structure is transferred by applying a drop of liquid PDMS to a coverslip and imprinting the topology using a passivated PDMS stamp on the uncured PDMS.

**Fig. 2.**
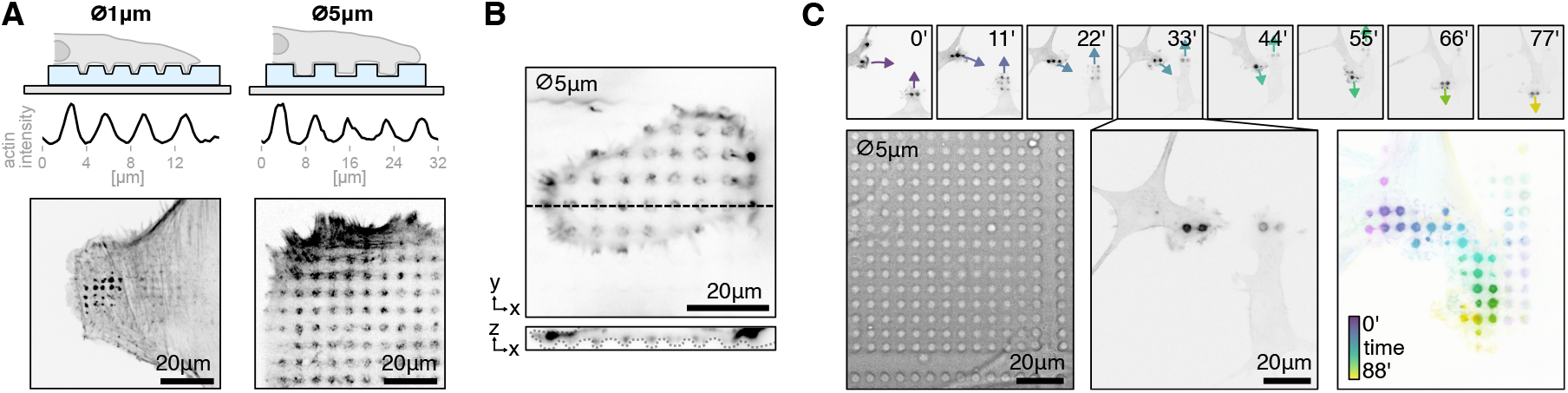
Microtopology: **A** Cells are seeded on a thin PDMS layer with pits or pillars with diameters of 1μm and 5μm. Fluorescence imaging and intensity profiles show the regular spacing of actin clusters forming at pit locations (confocal z-slice, more data fig. S6 and supp. movie M1). **B** 3D imaging shows that cells form actin rich protrusions into 5 μm pits (confocal z-stack). **C** Timelapse microscopy reveals rich actin dynamics of two cells migrating on 5 μm pillars. Small pictures show selected frames of the movie with arrows indicating the migration direction, brightfield image shows PDMS surface with 5 μm pillars. Last image shows temporal-color code of actin movie (supp. movie M2). All images in this figure are single confocal z-slices.

The coverslip is subsequently placed in an oven to harden the PDMS, after which the stamp is carefully removed, leaving behind a surface with well-defined pits or pillars. REF52 fibroblast cells expressing a fluorescent biomarker for F-actin [48], are then seeded onto the patterned surface.

Live cell imaging reveals that the cells form actin-rich protrusions extending into the pits, indicating an active response of the cell to the engineered surface topology (fig. 2B, supp. movie M1). Timelapse imaging results of cells on pillar structures are shown in fig. 2C and suppl. movie M2. Striking F-actin patches form at the leading edge of the cell, completely engulfing the pillars.

### Patterning of surface chemistry to control cytoskeletal shape or colony growth

Micropatterning enables the modeling of cell and tissue microenvironments by chemically patterning surfaces. This technique allows researchers to control cell and tissue morphology by enforcing specific shapes, facilitating causal investigations into the relationship between morphology/geometry and function, or reducing heterogeneity by standardizing cell shape [22, 29, 49].

The most widely used micropatterning method today is microcontact printing [50]. In this approach, extracellular matrix proteins such as fibronectin or laminin are stamped onto a glass slide using a PDMS stamp. The unstamped regions are then coated with PLL-g-PEG, a non-adhesive polymer that prevents cell attachment [50] (fig. 3A). Commercially available slides with experiment-ready, standardized patterns simplify this procedure and enhance reproducibility; however, custom patterns are often required depending on the experiment or the adhesive properties of specific cell lines.

**Fig. 3.**
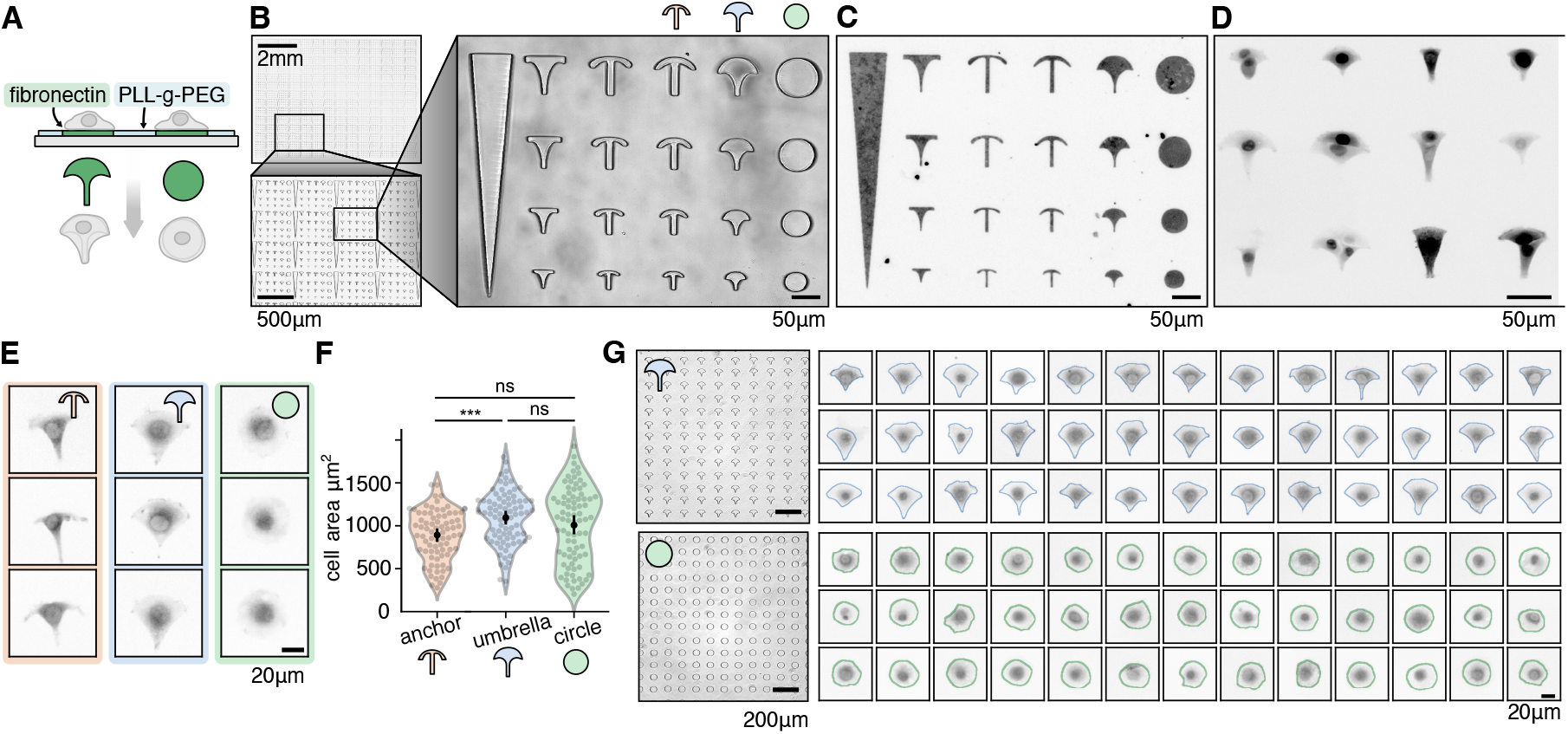
Micropatterning: **A** Glass is patterned with patches of matrix proteins like fibronectin which cells attach to, and PLL-g-PEG which is non-adhesive. The cells adopt the shape of the pattern. **B** Microfabricated PDMS stamp containing different designs and sizes to screen. **C** Fluorescent laminin showing precise surface patterning (full chip shown in S4). **D** Cells attaching to the micropatterns. **E** Sample cells on three different shapes (anchor, umbrella, circle). The cells on the anchor pattern are more contracted compared to the umbrella shape. **F** Quantification of cells on different shapes shows that cells on umbrella patterns are significantly larger than on anchor patterns (t=4.33, p<0.001, df=159). **G** The best patterns from the screen are selected to create large arrays of the same pattern. The morphology of hundreds of cells can be homogenized per experiment. Crops show cells with segmentation outline, automatically detected and filtered by morphological features. Brightfield images in B and G are high-pass filtered to reduce vignetting and out of focus artifacts.

Our method allows for the rapid prototyping of PDMS stamps with various designs. As a proof of concept, we designed variations of the commonly used anchor shape pattern (fig. 3B) to identify the optimal size and design that allows optimal spreading of NIH3T3 fibroblasts). Instead of stamping the fibronectin with a PDMS stamp, we first coated the whole slide with fibronectin (or fluorescent laminin for quality control) then protected areas by placing the PDMS stamp on top. We then etched away the unprotected areas, with a plasma cleaner. The etched regions are subsequently filled with PLL-g-PEG, ensuring that cells adhere exclusively to the fibronectin-coated regions while they can be washed away from the non-adhesive PLL-g-PEG areas. This left us with patterns with sharp edges, clearly reproducing fine structures of the shape down to a few μm (fig. 3C) across an area of a few mm (fig. S4, to which the cells attaches to well (fig. 3D). For single cell patterns, coating the entire slide first and then using the stamp to protect specific areas, rather than coating the stamp and stamping onto the glass, has been more convenient and reproducible for us.

Our goal was to identify patterns that accommodate a single cell while promoting optimal spreading without inducing excessive retractions or cell detachment. For example, many cells on the anchor shape seemed to collapse on the sides, while on the umbrella shape they were more spread out (fig. 3E). We automatically segmented the cells with Cellpose3 [51] and measured their area (anchor: 891.99^2^± 287.2 (n = 79), umbrella: 1094.29 μm^2^ ± 304.68 (n = 82), circle: 1007.31 μm^2^ ± 439.72 (n = 80)), and found that it is significantly higher in the umbrella shape versus the anchor shape (independent t-test: t = 4.33, p < 0.001, df = 159) (fig. 3F).

After screening different pattern variations, we selected the most suitable design and fabricated an array featuring the optimal shape and size for our experiments. This patterned grid enables the production of hundreds of cells with a standardized cytoskeletal organization, ensuring morphological uniformity (fig. 3G).

Additionally, we demonstrate the value of our approach at the tissue scale to generate circular patterns that are routinely used in the stem cell field to generate 2D gastruloids of defined diameters, similar to Warmflash et al. [49] (fig. 1B, S5). For stem cell colony formation with large diameters (250-1500 μm), a basement membrane extract is stamped onto a glass slide using a PDMS stamp, followed by cell seeding. No non-adhesive coating is needed.

### Multilayer microfluidic devices to study confined cell migration

Beyond classic 2D migration models, cells migrate in a 3D environment in vivo [52]. Microfluidic devices with precisely shaped constrictions have provided a way to study 3D migration under well-defined conditions. These devices have been used to investigate nuclear deformation [53] and the cytoskeletal mechanisms that generate the forces required for cells to squeeze and migrate through narrow gaps [10, 11].

We demonstrate that our fabrication method can produce such microfluidic chips with customizable geometries (fig. 4A-D). To enhance flow through the channels, we print larger layer heights for the regions leading to the constrictions. The device master mold is fabricated by iteratively spin-coating and exposing layers of 3D printing resin, aligning the layers precisely using a fiducial marker and the microscope camera. The fiducial marker is printed as part of the microfabricated structure (X-shape visible in top left corner fig. 1C). After calibrating with a single layer and printing two stacked layers, we observed a 4.92% error from the expected layer thickness, indicating that spin coating resin directly on glass or on a previously printed layer does not significantly affect spin coating properties (fig. S7). A UV-free light source is used during alignment to prevent accidental polymerization of the resin.

**Fig. 4.**
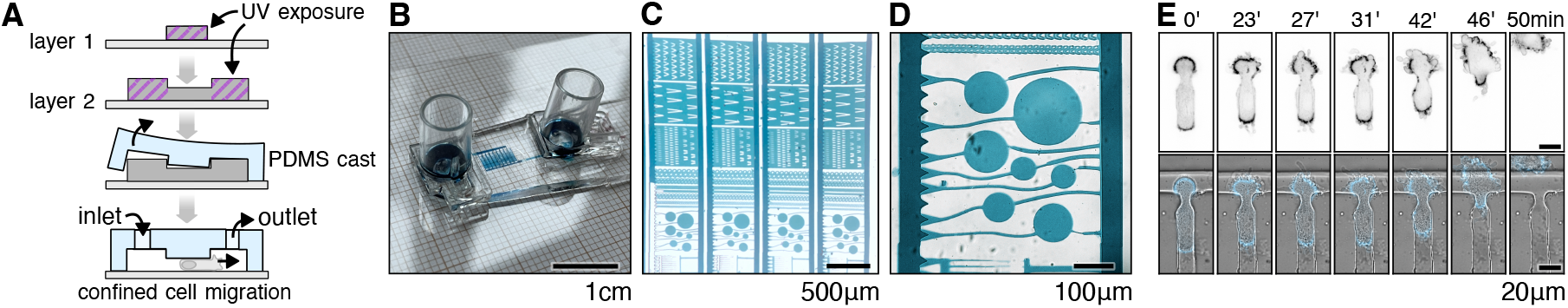
Microfluidics: **A** 2.5D structure is achieved by iteratively spin-coating and exposing 3D printing resin. Supply channels are printed with a larger layer height to increase fluid flow rate. The microfluidic device is built by punching inlet and outlet holes, then plasma-bonding the PDMS stamp onto a coverslide. **B** Complete microfluidic chip with adapters for syringes or automated pumping systems. Chip is filled with food coloring to visualize the channels. Grid lines for scale (small squares are 1 mm, large squares 1 cm) **C** Section of the chip photographed with a smartphone through a microscope ocular, manually adjusted white balance. **D** Brightfield image 20× objective. As the used microscope camera provides only grayscale images, multiple exposures with different filters are merged and color balance is adjusted to achieve an RGB image. **E** A different chip design, showing a timelapse movie of a cell migrating trough a constriction (supp. movie M3).

REF52 cells expressing a biosensor for F-actin are seeded onto the PDMS device passivated with PLL-g-PEG and are allowed to settle for two hours. Spinning-disk confocal imaging enables clear visualization of actin patterns along the cortex, as cells migrate through the constrictions (fig. 4, supp. movie M3).

### Imprinting agar chambers for long-term tracking of *C. elegans*

Regulating organ size during development is crucial, as even minor imbalances in growth rates can lead to significant deviations in organ proportions. Studies using *C. elegans* have demonstrated that organ size scaling remains remarkably consistent across individuals [54, 55]. To track growth over several days by imaging, individual worms were placed in agarose microchambers [8] (fig. 5A). While microfluidics-based systems allow temporary immobilization of worms for imaging weak fluorescent signals [56], the microchamber approach discussed here enables imaging of a larger number of animals in parallel, is simpler to manufacture, and does not require vacuum or pressure pumps.

**Fig. 5.**
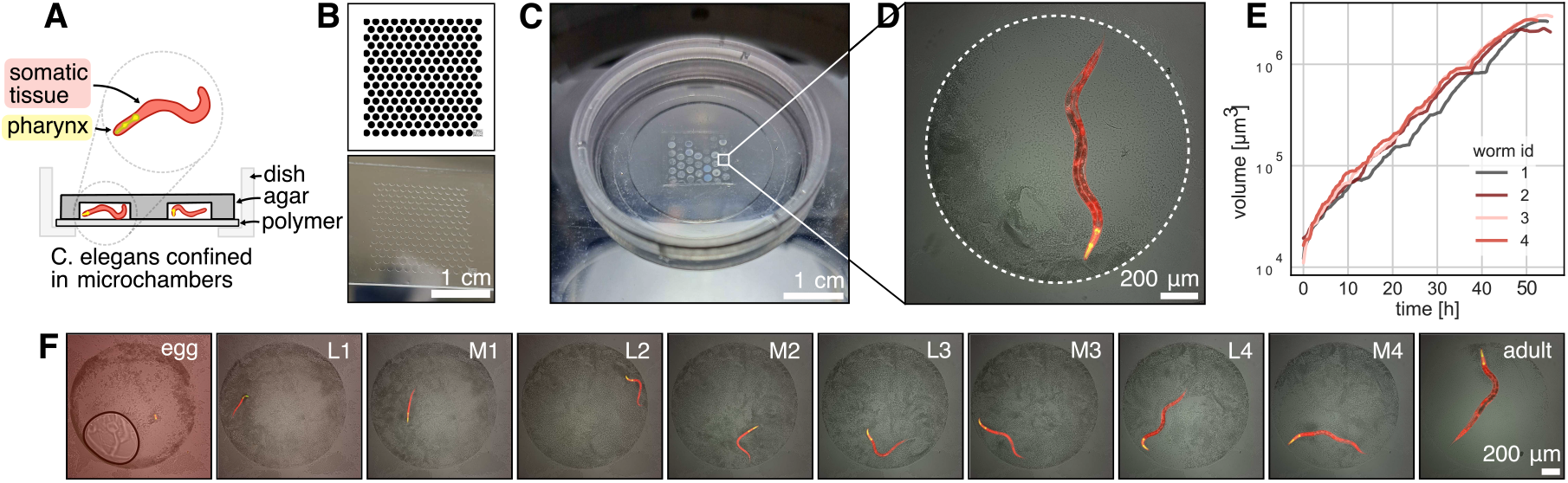
Microchambers. **A** Agar microchambers for long-term tracking of *C. elegans* tagged with fluorescent markers expressed in the pharync and somatic tissues. **B** Digital mask containing an array of microwells and photo of the corresponding structure made from 3D printing resin. From this structure a PDMS stamp is molded. **C** The PDMS stamp is used to imprint microwells into agar. Photo shows an array of microwells loaded with worms loaded in an ibidi dish for long term imaging. **D** Microscope image of a microchamber containing a worm. The brightfield channel shows the microchamber and bacteria as food source, fluorescent channels the pharynx (green, *myo-2p::GFP*) and somatic tissues (red, *eft-3p::mScarlet*). **E** The microchambers restrict the movement of the worms, allowing to track the individual growth over multiple days. **F** Timeseries showing the complete lifecycle of one worm (supp. movie M4).

Microchamber designs can be ordered from lithography companies. Multiple designs can be batched into a single order to reduce costs. Depending on the size of the design and the company, around 20 designs can be ordered in one go, resulting in an approximate cost of 60 C per design and a delivery time of a few weeks. Our method allows for customization and testing of different chamber patterns within a day. For instance, chamber size can be adjusted to match the microscope’s field of view, optimizing space and food quantity for each worm while preventing it from moving out of frame. Our method reduces material costs to <1 € per chip.

Here, we present circular microchambers arranged in a honeycomb pattern (fig. 5B), enabling long-term tracking of many individual worms in parallel. For preparing worm imaging chambers, a PDMS master is first fixed to a glass slide using double-sided tape and is plasma treated to clean it and enhance hydrophilicity. The master is then used to cast chambers by sliding it into melted agarose gel on a glass slide. After a brief curing period, the agarose is trimmed to retain only the wells and surrounding space.

Bacteria are added to the wells as a food source. Worm eggs are collected from culture plates and individually placed into the wells using an eyelash pick, ensuring only eggs at the 2-fold stage are selected. The agarose wells are then inverted onto an imaging dish with a gas-permeable polymer bottom (fig. 5C).

To seal the system, the dish is covered with low-melting agarose followed by a PDMS overlay, and parafilm to prevent evaporation. The PDMS cures at room temperature during image acquisition. A custom plate holder allows simultaneous imaging of six dishes on a single microscope. The worms are tagged with fluorescent proteins, marking the pharynx and somatic tissues (fig. 5D). Timelapse imaging and automated image analysis allows to track the growth of individual worms over multiple days (fig. 5E), capturing the complete lifecycle, from hatching to laying eggs (fig. 5F, supp. movie M4).

## Discussion

Photolithography and soft lithography are widely used in biological research for microfabrication due to their high spatial resolution and versatility. However, these methods require specialized equipment, trained personnel, and the use of toxic chemicals, which can be a barrier for many laboratories. In contrast, 3D printing offers a low-cost and accessible alternative but lacks the spatial resolution needed for many microfabrication applications.

In this work, we repurpose a fluorescence microscope designed for targeted photostimulation for microfabrication. By combining 3D printing with established lithography techniques, we achieve micrometer-scale precision over centimeter-scale areas while maintaining rapid prototyping capabilities. This approach makes microfabrication more accessible and applicable to a wide range of research questions.

We now approach microfabrication much like conventional 3D printing: as a low-friction tool for rapidly designing custom solutions. When an idea arises, we often ask: Could microfabrication be a solution here? The ease of use allows us to quickly fabricate a prototype, yielding results within a day. The low cost minimizes the risk of trying. This accessibility has led us to integrate microfabrication into numerous projects within the lab and institute, where we previously wouldn’t have considered it, simply because it is now so straightforward to implement.

Previous efforts to simplify microfabrication workflows have focused on eliminating the need for clean room facilities [32], using glass [30] or polyethylene terephthalate (PET) [57] instead of silicon wafers as a substrate, and finding alternatives to the expensive photoresist SU-8 [18, 58]. Maskless lithography systems utilizing scanning stages [59, 60] or projectors [18, 30, 33–35, 61] have been introduced to increase iteration speed. While many maskless systems rely on custom hardware setups, some labs have leveraged commercial DMD systems and microscopes [18, 35]. Another group has developed a method to facilitate the separation of PDMS structures without using chlorosilane coating [62], a widely used but highly toxic chemical that releases hydrochloric acid upon contact with water. Our approach integrates these scattered simplifications into a single workflow and builds upon them to further streamline the process.

Our microfabrication workflow only requires 3 non-standard consumables: TMSPMA, PDMS and 3D printing resin (all listed in table 1). The use of 3D printing resin as a substitute for SU-8 eliminates time-sensitive baking steps and reduces the need for extensive glass slide cleaning. Instead of using highly corrosive piranha solution, a short submersion in TMSPMA solution is sufficient to ensure strong bonding between the printed structures and the glass substrate. Leveraging microscope control software with Python scripting allows for seamless customization and automation of the printing process, which is particularly beneficial for iterative design processes.

Using μManager, our method is compatible with a wide range of existing microscope hardware, ensuring reproducibility across different lab settings. While this paper focuses on DMD-based patterning, we have also explored the use of a total internal reflection fluorescence (TIRF) and fluorescence recovery after photobleaching (FRAP) system (iLas 2, GATACA) with galvo mirrors. Although this system offered considerably slower scanning speeds per field of view, it was still viable for producing useful micro-fabricated structures.

For labs looking for a dedicated microfabrication microscope, there are complete hardware solutions available commercially (Primo Optical Platform, Alvéole) that should be compatible with the methods presented in this paper.

Certain limitations remain: Plasma cleaners and DMD systems (see table 2 for required hardware devices) and can be expensive if not already available in the lab, although DIY alternatives have been demonstrated at a fraction of the cost [63, 64] of commercial systems. Among the required materials, PDMS is the most expensive ($162 / 0.5 kg), but the cost per fabricated device is low and the chips can be reused multiple times.

Looking ahead, future efforts will focus on developing lower-cost hardware solutions to further democratize access to microfabrication techniques. Digital light processing (DLP) printers contain much of the necessary hardware (UV lamp, driver board, DMD) at a fraction of the cost (Anycubic Photon Ultra is $300). The maker community has already made significant progress in adopting such systems for microfabrication [65]. Integrating such components into low-cost modular microscope systems, such as UC2 [66], could provide an affordable alternative to high-end microscope projection photolithography setups.

Our Python-based workflow, along with interactive fabrication features that allow regions of interest to be selectively exposed in real time using live camera feedback, presents opportunities for intelligent automation. Potential applications include automatic alignment and exposure compensation. Closed-loop positioning with computer vision could compensate for low-precision x/y stages by digitally adjusting the stimulation mask to correct mechanical stage positioning errors. By mixing dark particles into the 3D printing resin, we can generate unique speckle patterns that serve as fiducial markers for image registration, which in the future might enable high-precision microfabrication despite stage limitations. Real-time feedback capabilities could also enable the fabrication of UV-crosslinkable, cell-compatible hydrogels, allowing researchers to print structures in situ and capture cells of interest with high precision.

We hope this paper stands out by demonstrating the versatility of the technique across diverse biological applications. We believe the simplicity and versatility of our method will encourage broader adoption across the scientific community. Our detailed methods section, which includes practical troubleshooting tips, should further facilitate reproducibility. The open-source nature of the project is expected to inspire further developments and novel applications across diverse fields.

## Code availablability

The software is open source (BSD-3) and hosted on GitHub: https://github.com/hinderling/fabscope.

## ACKNOWLEDGEMENTS

This work has been supported by Uniscientia fellowship 187-2021 to OP, SNF Sinergia grant CRSII5_183550 to OP, SNF grant 310030_185376 to OP, Experiment (experiment.com) grant 10.18258/50706 to LH and AL, SNSF Eccellenza grant (PCEFP3_181204) to BDT, SNSF project grant (310030_207475) to BDT. LH received support through the Open Round funding for Young Scientists by the Faculty of Science of the University of Bern (2022-4). Microscopy experiments were performed on equipment supported by the Microscopy Imaging Center (MIC), University of Bern, Switzerland. We thank Samir Gupta (MIC) for his help with light power measurements.

## AUTHOR CONTRIBUTIONS

LH conceptualized the work. All authors contributed to the development of the method. LH, JF, RH, MK, BG, and SP acquired data. Figures were created by LH. LH and OP wrote the manuscript and acquired funding. All authors read and approved the final manuscript.

## Methods

### Microfabrication protocol

All of the microfabrication applications start by creating a 3D printed mold from UV-curable resin, from which a PDMS cast is made. Standard microscope slides are used as a carrier for the microfabricated structure for their sturdiness and availability.

#### 1. Prepare the glass slide for good adhesion

Clean the glass slide for 2 min in the plasma cleaner (other methods proposed in the literature are ultrasonic cleaning, cooking in deionized water, EtOH bath but led to worse results for us.)

Coat the slide with 3-(trimethoxysily)propyl methacrylate (TMSPMA) by submerging it in a 2% solution of TMSPMA (solvent: 96% EtOh, 4% Keton) for 5 minutes. After incubation the slide is washed in a EtOH bath for 5 min, then dried in the oven for 10 minutes at 70°C. To conserve TMSPMA solution, a small amount can be applied with a pipette instead of full submersion; due to the glass’s hydrophilicity from plasma activation, the solution spreads evenly across the surface.

#### 2. Coat the glass slide with the UV-curable resin

The thickness of this layer determines the final height of the structure. The thickness is adjusted by varying the rpm of the spin coater and is dependent on the viscosity of the resin. The viscosity differs strongly between brands, and also increases as solvents from the resin evaporate. To minimize this, we recommend to prepare aliquots of resin in 50 ml falcon tubes instead of opening the storage bottle many times. We first deposit ~1 ml of resin onto the glass slide, a pipet can be used with a cut-off tip to facilitate flow of the viscous fluid. Be careful to not create any bubbles, as they result in uneven layer height. The resin is roughly distributed on the slide by tilting in in different directions before spinning. We recommend a two-step program for the spin coater, a slow first step with 1/2 of the final speed or 800 rpm (whichever is lower), for 10 seconds to distribute the resin evenly on the slide, then a second step for 20 seconds with the final speed to achieve the required thickness. Spinning with high rpm directly may result in the resin being slung off the glass slide. The slide is now ready for UV exposure. When carrying the slide between rooms, cover it to reduce contamination with dust particles.

#### 3. Expose the pattern with UV light

The slide is placed into the microscope, resin side towards the objective. For inverted microscopes, make sure there is no excess resin that could drip into the objective. Find the focus plane, load the pattern and set the exposure time (described in detail below).

#### 4. Remove uncured resin

Immediately after pattern exposure, the uncured resin is washed by placing the slide in an Isopropanol bath for 5-15 minutes, depending on layer thickness. Any remaining uncured resin, e.g. in narrow gaps, can be washed out efficiently by spraying Isopropanol from a pressure sprayer. When producing many slides, to reduce Isopropanol waste, multiple baths can be used, moving the slides sequentially from the dirtiest to the cleanliest. After washing, the glass and 3D print should be clear and not show any smears or diffuse residues.

#### 5. Post-curing the resin

The resin is exposed under a homemade UV-light (see *Calibrating the UV-exposure time* below) for 5 min to activate all photoinitiators remaining in the resin and complete hardening of the pattern. To evaporate all volatile species that could cause curing inhibition (see below), keep the slide at 70°C over night.

#### 6. Creating a PDMS replicate

The 3D printed structure is used as a mold to cast a negative PDMS mold (see below for PDMS preparation). To improve separation between the PDMS and 3D print, the 3D print is plasma activated for 30 s, then dipped into methanol bath. The PDMS coated with methanol is left at room temperature (5 min), then the remaining methanol is discarded. The slide is left to dry at room temperature an then placed in the oven at 70°C to complete the drying process. Now the PDMS can be poured onto the 3D print. Slowly pouring from one side avoids trapping pockets of air. To remove any air bubbles, the slide can be placed in a vacuum pump for 30 min or until no more bubbling is visible. The slide with the PDMS is then placed into the oven at 70°C for around 2 hours or until the PDMS is hardened and non-sticky to the touch. Now the PDMS mold can be peeled of the glass slide with the 3D printed structure. If there are issues with adhesion between the 3D print and glass, the 3D printed structures can detach during this step and stick to the PDMS instead (check below fora trouble shooting guide).

#### 7. Inverting the PDMS replicate

We recommend to use the negative mold as template to create any PDMS structures actually used in experiments, as they show greater durability than the 3D printed slide for repeated mold use (resistance to accidental scratches, ease of separation). To create a positive PDMS structure from the negative mold, the negative mold has to be passivated otherwise the PDMS structures are not separable. Similar to the passivation of the 3D print, the negative mold is plasma activated for 10 s, then coated with methanol [62], which is left for 5 minutes then placed in the oven for drying at 70°C. Now the PDMS can be poured, again taking care to avoid creating bubbles and remove them if necessary by placing the mold into a vacuum unit.

##### PDMS preparation

PDMS is mixed in a 1:10 ratio of crosslinker and elastomer in a large batch, then aliquoted into 50 ml falcon tubes and centrifuged to remove trapped air bubbles. Centrifugation works faster and produces less waste than removing air bubbles with the vacuum pump. Aliquots can be stored at −20°C for months without curing. When using the PDMS, the desired quantity can be poured cold directly from the cold falcon tube.

##### Finding the focus plane

The slide is placed into the microscope, resin side towards the objective. For inverted microscopes, make sure there is no excess resin that could drip into the objective. To find the focus plane, a checkerboard is projected onto the glass slide with red light. Moving in the z-axis towards the slide, the first focus plane is resin surface, the second focus plane corresponds to the surface of the glass slide. We got best results focusing on the surface layer of the resin (1st focus plane) or slightly below.

##### Calibrating the UV-exposure time

Set the correct UV-light exposure time, which is dependent on the brand of resin, light source intensity, DMD, objective, and even the design of print. Very small or thin patterns may require increasing the exposure times. The multitude of factors makes calculating the exposure time difficult, and its easiest to just perform an exposures test-series. The test series can be performed automatically using code we provide. The quality can be inspected directly after cleaning with isopropanol, so the whole calibration procedure should not take more than 10 min. For our setup, at a layer thickness of 14 μm, we use 600 ms for 4x magnification objectives, 150 ms for 10×, 30 ms for 20×. We use a LED light source (Lumencor Spectra X) at 395 nm with a 395/25x excitation filter (Lumencor) and measure 2.96 mW at the 10× objective surface. The fraction of activated DMD pixels is linear to the power reaching the sample (see fig. S8, linear fit of the data shows a R^2^ value of 1.00, slope 2.71 mW/100% ON pixels, intercept −0.02 mW).

##### Avoiding curing inhibition

One of the main difficulties when using 3D printing resins with PDMS is that unactivated photoinitiators in the resin can leach into the PDMS and prevent it from polymerizing, while remaining resin monomers can increase adhesion of PDMS to the 3D printed mold, making them difficult to separate, see the study by Venzac et al. [39]. They find that 11 out of 16 tested resins can be treated using a combination of 120°C heat treatment and UV exposure with a total curing time below 135 min, which should be considered if protocol duration is essential. For us curing at 70°C over night was a practical tradeoff, also preventing issues warping/detachment of the 3D print at high temperatures. We also have one oven running at 70°C in the lab anyways, using it instead of running an additional oven at 120°C is also more energy efficient. We built a UV-curing station by cladding a box with reflective aluminium foil and using a 405 nm LED unit (CR-6565-4CLED) marketed as replacement part for Epson flatbed printers. It comes packaged with an aluminum heatsink and cooling fan. Available online for < 20USD with different wavelengths (365nm, 385nm, 359nm, 405nm). The unit runs at 24V / 40W, we use a AC85-265V to DC24-36V converter (LED driver YJ-TG50W-1300) as power supply.

##### Plasma cleaning and activation

We perform all our plasma cleaning or activation steps with atmospheric gas mix at 3 torr pressure, only varying the timing. PDMS can start to break down if activating for too long, leading to a rough surface texture that looses its adhesive properties. This can be observed visually as PDMS appearance of an opaque surface layer.

##### Trouble shooting resin-glass adhesion

The composition of glass slides used can make a difference, which is difficult to figure out as not all brands are clear about the additives and surface coatings used. We had some experiments fail when using a freshly opened package of a very old stock of slides, which upon close visual inspection showed oily smeared surface. TMSPMA coating was integral to ensuring good adhesion. We tried coating the slides in batch and storing for later experiments, but they develop a visible layer of impurities if stored after some weeks which decreases adhesiveness. For best results use freshly coated slides.

##### Trouble shooting PDMS separation

For separation of PDMS and glass/resin, it is important to plasma activate the slide long enough. 1min 30 sec provided good separation results, while anything below 30 seconds lead to difficulties separating the glass and PDMS. Duration of plasma activation is also important for PDMS-PDMS separation. While we get good results with 10 s activation time, layers bonded much too strongly when activating 30 s or longer. Methanol incubation time did not seem to affect separation quality in either of the procedures.

##### 3D imaging of the PDMS stamps for quality control

Liquid fluorescent dye under UV light is extracted from a yellow highlighter pen by breaking it open and squeezing the ink from the fibers into a solvent (water or ethanol). The negative PDMS stamp is plasma-activated and coated with the extracted fluorescein. It is then pressed against a coverslip, allowing excess fluorescein to be expelled from the sides. Z-stacks are acquired using a confocal microscope. Alternatively, a positive stamp can be pressed onto a coverslide, and the empty volume can be filled with fluorescein from the sides (florescence exclusion). We used the second method to measure z-layer height accuracy. We printed test patterns at different rpm (200, 400, 800, 1600, 3200), and replicated the structure in PDMS with double casting. Brightfield and fluorescence exclusion z-stacks of a sample pattern (400 rpm) are shown in supp. fig. S3.

### Micropatterning for single cell fibroblasts

#### 1. Preparing the well plate

A 24-well plate is plasma activated for 30 seconds, then coated with 250 μl of 10 μg/ml fibronectin (human plasma fibronectin purified protein, Merck) diluted in MilliQ water. Plasma activation increases the hydrophilicity of the glass, decreasing the volume of fibronectin solution required to coat the surface. The plate is incubated for 1 hour at 37°C or overnight at 4°C. It is then washed thoroughly with PBS and stored at 4°C. Before use, PBS is aspirated, and the wells are allowed to dry in the hood. The PDMS stamp is placed onto the coated wells. The well plate is plasma activated for 1 minute and 30 seconds. After activation, the PDMS stamps are removed, and the wells are washed with MilliQ water. A solution of 250 μl PLL-PEG (100 μg/ml) (PLL(20)-g[3.5]-PEG(5), SuSoS) or F-Pluronic (5%) (Pluronic F127, Bioreagent) is added and incubated at room temperature for 1 hour. F-Pluronic is cheaper but has decreased anti-adhesive properties for some cell lines. The solution is aspirated, followed by washing with PBS. The wells are covered with PBS and are ready to use. Alternatively, they can be stored at 4°C for months.

#### 2. Seeding cells

PBS is aspirated from the wells, and 300 μl of growth medium is added. ~20’000 cells are then seeded per well into growth medium, and are incubated at 37°C and left to attach to the micropatterns for 3 h (varies depending on cell line, pattern, and cell density). Seeding density is checked, and the cells are gently washed with PBS until the desired density is achieved.

##### Reusing the PDMS stamp

The PDMS stamps can be cleaned with ethanol and re-used after drying. If soaked in ethanol for long, the stamp can acquire a opaque texture, which disappears after drying. Drying can be sped up by placing the stamp into a 70°C oven.

##### Quality control micropatterns

The well plate is coated with 1 μg/ml fluorescent laminin (LMN01-A, Cytoskeleton) for 1 hour at 37°C or overnight at 4°C. It is then washed with MilliQ water and covered in MilliQ for storage at 4°C. Before stamping, MilliQ is aspirated, and the well plate is dried in the hood. The pattern is stamped using the plasma etching technique described above, and the fluorescent pattern is imaged.

### Micropatterning for stem cell 2D gastruloid

#### 1. Preparing the PDMS stamps

Stamps are cut to appropriate size, so they can easily fit the well of a 24-well plate. Sterilize stamps in Ethanol and then dry them under the hood.

#### 2. Preparing the wellplate

Coat a 24 well plate with (3-Mercaptopropyl)trimethyloxysilane by putting the plate open in the dessicator with 100 ul of (3-Mercaptopropyl)trimethyloxysilane in a falcontube lid. Pump is activated for 5 min and then turned off, stamps are left for another 30 min in the dessicator, then put in a 80°C oven.

#### 3. Stamping matrix

1:100 Geltrex solution (~50-100 μl per stamp) is added to dried stamps and put in the incubator for at least 30 min. Extra Geltrex is removed and stamps are left to dry (can be observed under the microscope). Once stamps are mostly dry, they are picked with up with tweezers sterilized in ethanol. Stamp is flipped and placed it in the center of a well with one motion. Stamp is gently pressed. This is repeated for all stamps. Stamps are then left for 20 min before being removed with tweezers with one motion. Wells are rinsed well with PBS.

#### 4. Coating non-adhesive polymer

400 μl of 5% Pluronic-F 127 is added to each well for 1 h. Each well is then gently washed 3x for 5-10min in PBS. Plate is now ready to use.

#### 5. Seeding cells

1ml of StemPro Accutase is added to the cells to dissociate. Cells are put back in the incubator and regularly checked. When most cells are dissociated (after 3-5 min), they are thoroughly pipetted and put in a 15 ml falcon tube. 7 ml of E8 flex (A2858501, ThermoFisher) is added, then the tube is centrifuge at 1000rpm for 1min. Cells are resuspended in 1 ml E8 flex with 10uM ROCK inhibitor (A3008, Apex Bio Lubio). Cells are counted, ~400’000 cells are seeded per well in 400 ul E8 flex with 10 uM ROCK inhibitor. When cells adhere, change to E8 flex medium without ROCK inhibitor (after 2 h or next day).

### Microfluidic chip

#### 1. Preparing the PDMS Stamp

The PDMS stamp containing the microfluidic channels is placed channel-side up on a cutting mat or a piece of cardboard. Access holes are created by punching through the chip using a biopsy punch, allowing connection to the channels from the outside.

#### 2. Bonding the PDMS stamp to the glass slide

The PDMS stamp and a large coverslip (Matsuinami Micro Cover Glass 50×70 mm, 0.13~0.17 μm)are placed channel-side up in a plasma oven and activated for 30 seconds. After activation, the PDMS stamp is gently placed onto the glass slide. By applying light pressure, the surfaces are brought into contact. Flipping the chip and reflecting uniform light from below helps identify bonded and non-bonded areas—bonded areas appear as dark patches, while non-bonded areas reflect more light. Localized pressure can be applied using a finger or a pen to ensure full bonding; however, excessive force may collapse the channels, causing the ceiling to irreversibly bond to the glass bottom. Once all areas are bonded, the chip is placed in an oven at 70°C for 15 minutes to complete the covalent bonding process.

#### 3. Adding fittings to inlets and outlets

Fittings are lightly dipped into a small puddle of PDMS. Excess PDMS should be removed by gently pressing the stamp onto a piece of paper to prevent clogging of the punched channels. The fitting is then positioned over the pre-punched inlet or outlet hole. The liquid PDMS should form a continuous seal around the entire circumference of the interface between the adapter and the PDMS stamp. The chip is then placed back into the oven to ensure a strong bond between the chip and the fittings. Additional PDMS can be applied externally to reinforce the bond if needed.

#### 4. Preparing the chip for the experiment

The microfluidic chip is plasma-activated for 1 minute to render the channels hydrophilic, facilitating the flushing process with medium. For the experiments presented in this paper, the channels are passivated with a non-adhesive coating by flushing the chip with a 0.5 mg/ml solution of PLL-g-PEG in PBS, followed by incubation at 37°C for 30 minutes. The PBS is then replaced with the experimental medium by adding it to the inlet port. Negative pressure is applied at the outlet using a syringe, drawing out the PBS until it is completely replaced by the medium. The chip is now ready for cell seeding.

#### 5. Seeding the cells

REF52 cells are washed twice with PBS and then detached using trypsin. The detached cells are transferred to a Falcon tube, and 5 ml of medium is added before centrifugation at 1000 rpm for 4 minutes. After centrifugation, the supernatant is removed, and 200 μL of fresh medium is added to achieve a high cell density. A drop of this dense cell suspension is placed in the chip inlet. A syringe is attached to the outlet fitting, and under microscopic observation, negative pressure is gently applied by pulling the syringe to draw cells into the channels. Once the desired cell density within the channels is achieved, the syringe is removed, and the chip is ready for imaging.

To validate the layer height accuracy when stacking multiple layers, we use the 3D imaging procedure for quality control described above. We assume the measured film thickness follows the well-established relationship that *h* is inversely proportional to the square root of the spinning speed *RPM* :

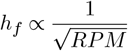

We print a circular pillar, one slide with one resin layer at 120 rpm and one slide with two iteratively printed layers at 210 rpm, aligning the pillar to stack its height. We plot a z-projection across the well border and measure the well depth using Fiji (fig. S7). We measure a depth of 43.93 μm for the single layer, and a total depth of 69.77 μm for the dual layer (34.885 μm per layer). For each measurement (*h*_*i*_, *RPM*_*i*_), we compute the constant:

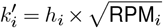

If the relationship holds, 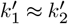. We calculate a final *error* = 4.93% using:

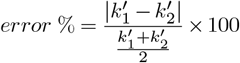

The inverse square root relationship between spin speed and film thickness has proven useful in calibrating our setup, enabling us to achieve the desired layer thicknesses in a single attempt, provided the RPMs are not too far off, as discussed in the results section.

### Agar microchambers for *C. elegans*

Method to create agar microchambers was previously described in Stojanovski et al. [55].

#### 1. Preparing the PDMS stamp

The PDMS stamp is fixed to a standard glass slide using double sided tape, then plasma activated for 30-60 seconds to clean it and make it hydrophilic.

#### 2. Creating the microchambers

Chambers are cast by sliding the PDMS stamp into a pool of molten 4.5% agarose gel dissolved in S-basal on a glass slide. After curing for 2 minutes, the sides of the agarose are cut to be left only with the wells and some space on the side. As a food source, the bacterial strain OP50-1 was grown on NGM plates by standard methods, scraped off using a piece of 3% NGM agar without cholesterol and then filled into the wells of the agarose gel.

#### 3. Placing the worms

Eggs are picked from plates previously grown and delivered to the spaces left on the side of the well array. Using an eyelash pick, wells are individually filled with eggs at 2-fold stage (younger eggs will not hatch).

#### 4. Imaging

Wells are inverted onto a dish of 3.5 cm diameter with a high optical quality gas-permeable polymer bottom (ibidi). The remaining surface of the dish gets covered with 3% low melting temperature agarose dissolved in S-basal, cooled down to below 42°C prior to application. The agarose gets topped with ~0.5 ml PDMS and the dish is sealed with parafilm to minimize water evaporation. PDMS is allowed to cure at room temperature on the microscope during the acquisition. Using a custom-made plate holder, six dishes can be imaged simultaneously on one microscope.

Analysis of microchamber experiments was done using a custom-made modular pipelining tool and an associated python package. Both are open source (BSD-3) and hosted on Github (https://github.com/spsalmon/towbintools_pipeline, https://github.com/spsalmon/towbintools)

## Supplementary figures

**S1** UV exposure calibration

**S2** X/Y resolution test using checkerboard pattern

**S3** Z-layer height calibration

**S4** Micropatterning of large areas

**S5** Micropatterning of 2D gastruloid

**S6** Actin clusters on microtopology

**S7** 2.5D printing

**S8** Measuring DMD output power

## Supplementary movies

**M1**`M1_pits_migration_1um.mp4`

Timelapse movie of a cell migrating on microtopology with 1 μm sized pits (shown in fig. 2A)

**M2**`M2_pillars_migration_5um.mp4`

Timelapse movie of cells migrating on microtopology with 5 μm sized pillars(shown in fig. 2C)

**M3**`M3_constricted_migration.mp4`

Cell migrating trough a narrow constriction of a microfluidic chip (shown if fig. 4F.)

**M4**`M4_C_elegans_microchamber.mp4`

Timeseries of a fluorescently tagged *C. elegans* worm confined in a microchamber. Complete lifecycle, from egg to adult (shown in fig. 5F).

**Fig. S1.**
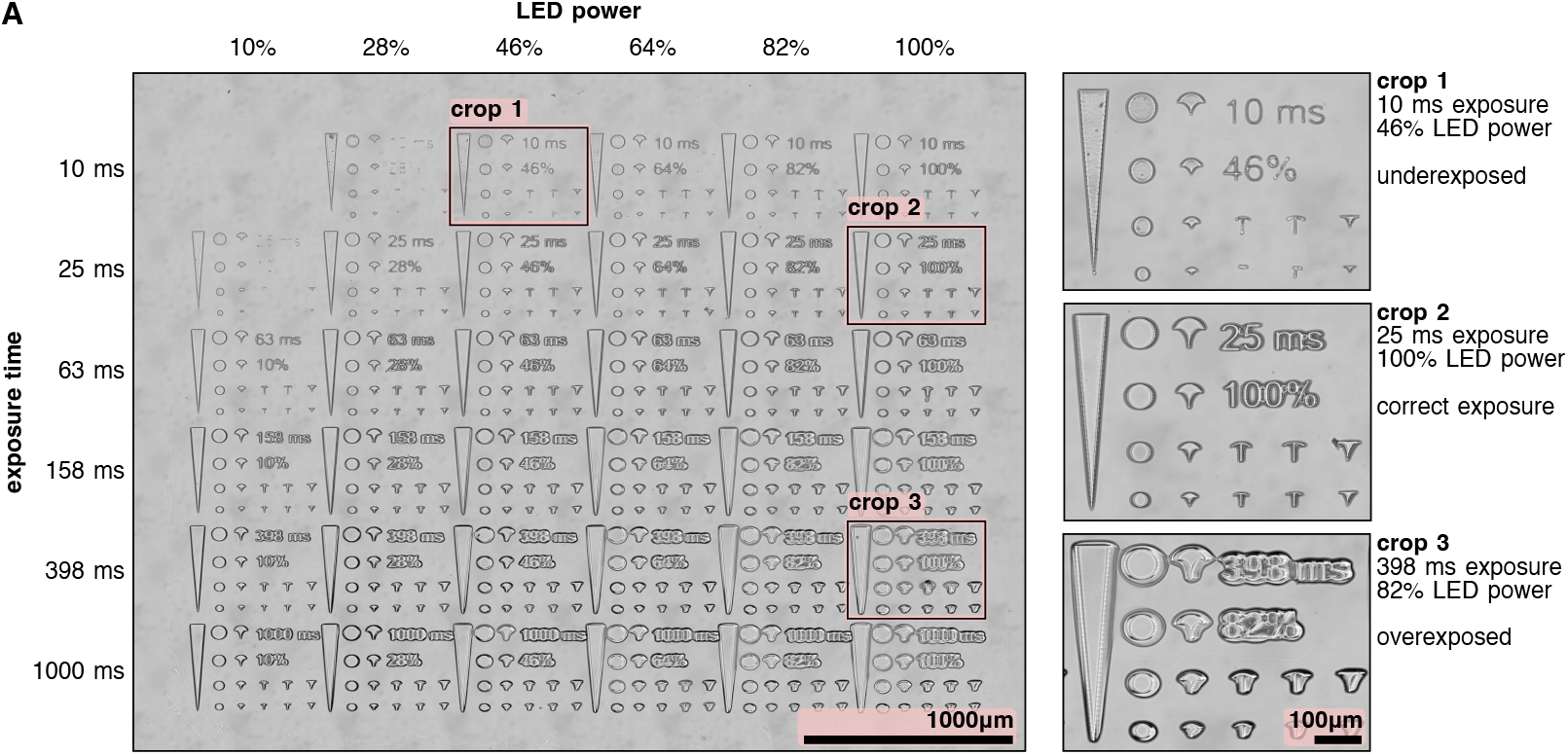
UV exposure calibration: Empirical grid search for exposure time and led power parameters. Different settings can be tested automatically in a single step, simplifying the calibration procedure compared to classical approaches. This test shows linearly spaced LED power settings, from 10% to 100% and logarithmically spaced exposure times, from 10 ms to 1000 ms. Duration of the test (exposing, washing, inspection) was under 5 min. Three crops show underexposed, correct exposure and overexposed FOVs. Slide is spin coated at 3000 rpm.

**Fig. S2.**
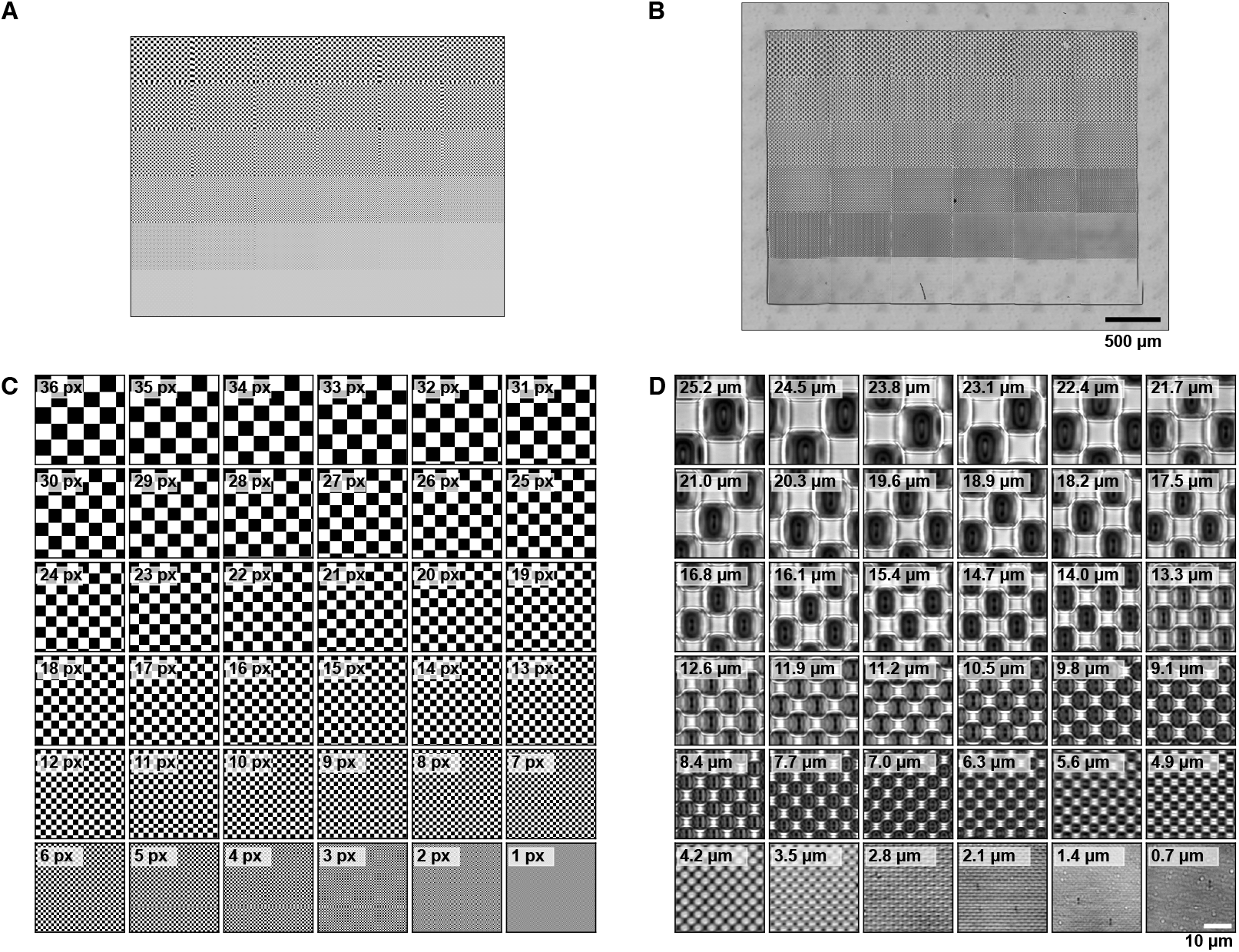
X/Y resolution test using checkerboard pattern. **A** 36 computer-designed test masks, containing checker-board patterns with smaller and smaller squares. Size per square ranges from 36 px in top left corner down to 1 px in bottom right corner. Each of the test masks has a resolution of 800×600 px which corresponds to the resolution of the DMD - one pixel in the mask maps to one pixel of the DMD. **B** Mosaic image of the printed test masks. The masks were printed with a 20× objective on slide spin coated at 3000 rpm. **C** Crop in on the different test patterns of the mask and **D** printed structure. Smallest pattern with clear edges has 4.9 μm pitch. For the checkerboard pattern, the effective resolution we can achieve with the DMD (4.9 μm or 7 px) is smaller than the theoretical resolution (0.7 m or 1 px).

**Fig. S3.**
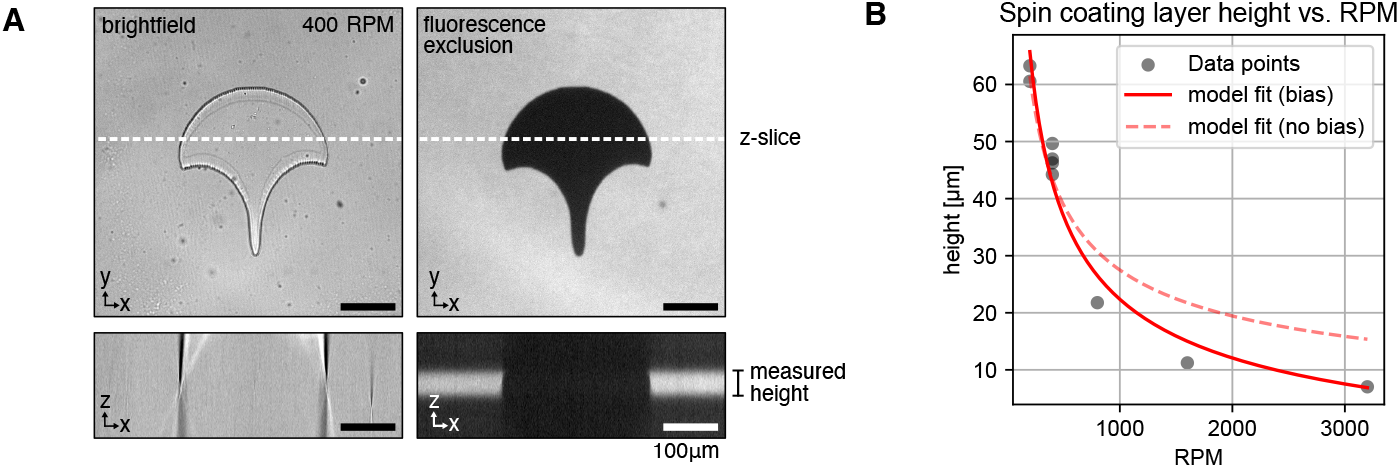
Z-layer height calibration: **A** Sample FOV showing brightfield and fluorescence exclusion confocal z-stacks. Height is measured manually using an x/z projection in ImageJ. **B** Plot showing the layer heights measured for different rpm (200, 400, 800, 1600, 3200). Two models are shown, with constant bias: 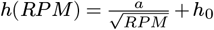 without bias: 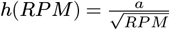. Model bias: *a* = 1110.09, *b* = −12.74 MAE: 3.60, RMSE: 4.07, R^2^: 0.96. Model no bias: *a* = 869.13, MAE: 4.86, RMSE: 5.99, R^2^: 0.90.

**Fig. S4.**
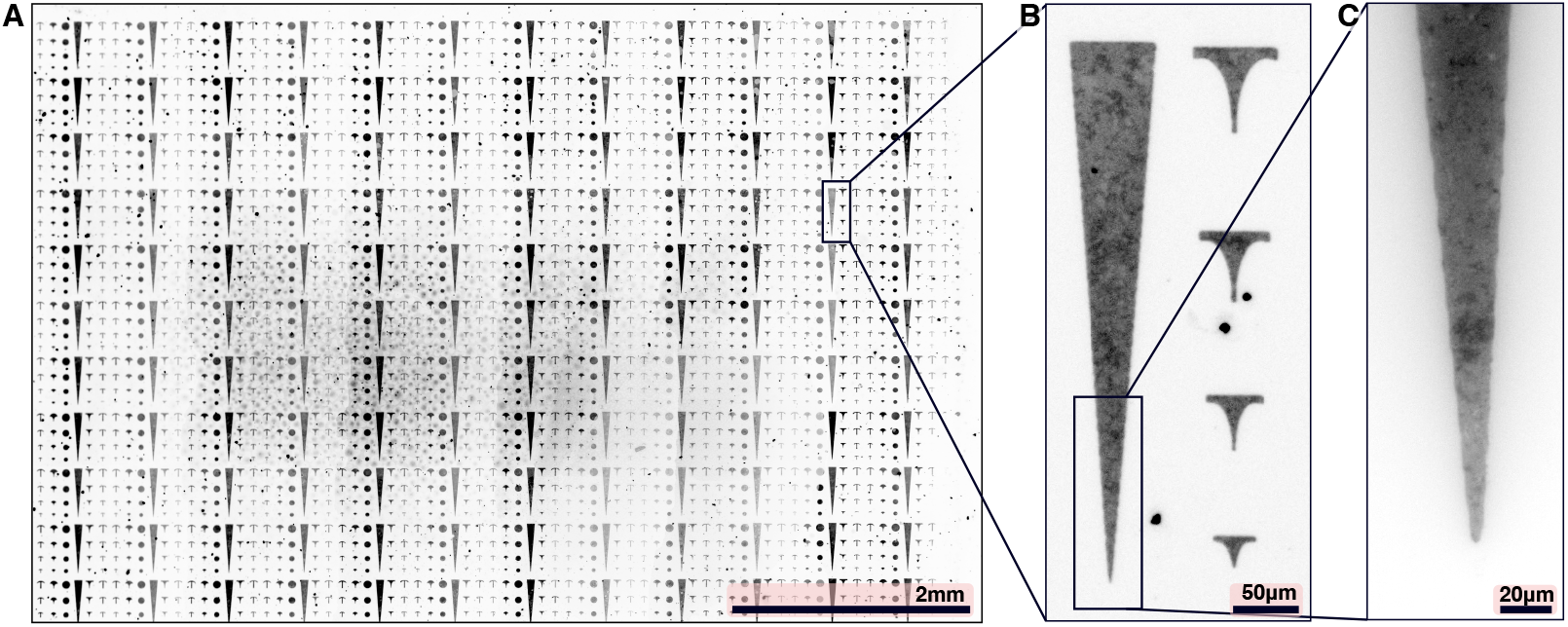
Micropatterning of large areas: Quality control of the micropatterning workflow using fluorescent laminin. **A**: Mosaic image (10× objective), some over-etching can be observed at the right edge, and some under-etching can be seen in the center of the slide. Different intensities of the patterns are artifacts of the automated image stitching. **B**: Close-up (10× objective). **C**: Close-up of triangle tip (40× objective).

**Fig. S5.**
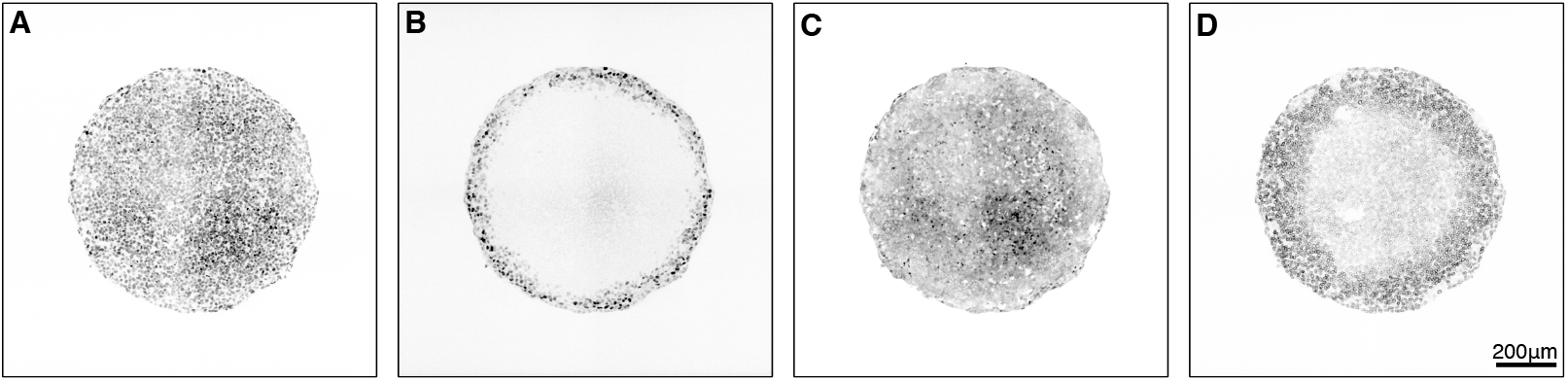
Micropatterning of 2D gastruloid: 800 μm disk pattern is used to create a circular gastruloid from human embryonic E9 stem cells. This figure shows single channels of composite in fig. 1B. **A** *H2B::miRFP670* nuclear marker **B** Brachyury immunostain pluripotency marker **C** *ERK-KTR::mClover* **D** *OCT4::POM121::tdTomato*

**Fig. S6.**
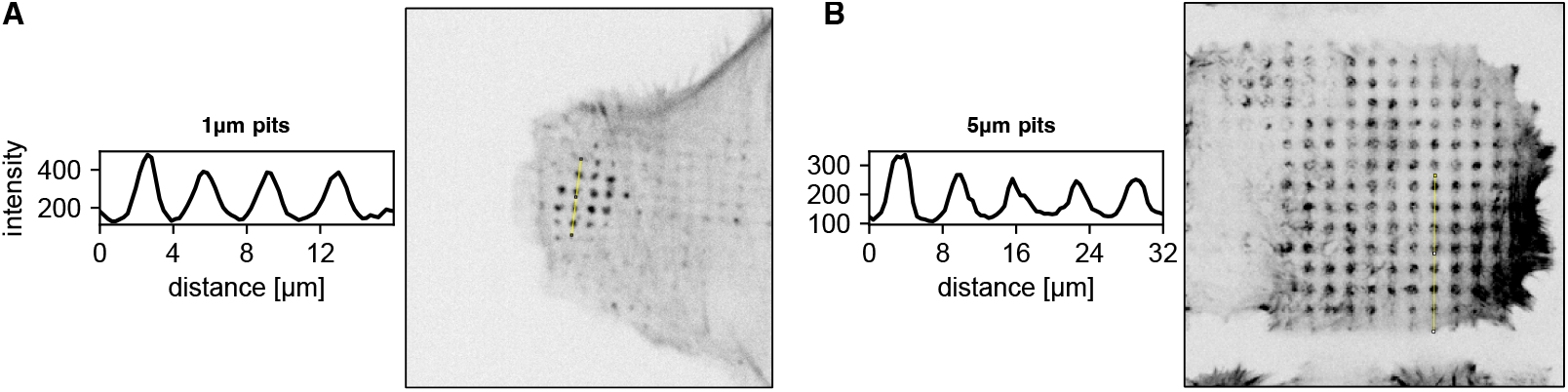
Actin clusters on microtopology: Lifeact intensity profiles of cells plated on **A** 1 μm sized pits and **B** 1 μm sized pits. Yellow lines indicate location of profile in image.

**Fig. S7.**
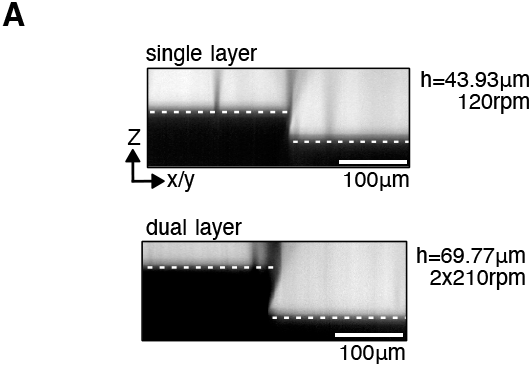
2.5D printing: Fluorescence exclusion z-slice of single and dual stacked layers acquire with a confocal microscope.

**Fig. S8.**
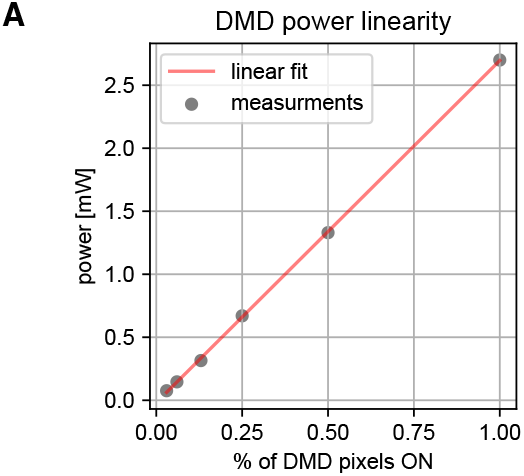
Measuring DMD output power: Datapoints show light intensity for different percentages of DMD mirrors set to ON state (1/32, 1/16, 1/8, 1/4, 1/2, 1/1). Energy is measured with a light meter held against a 10× objective. A linear fit of the data shows a R^2^ value of 1.00 (slope 2.71 mW/100%, intercept −0.02 mW).

## Footnotes

1 https://github.com/hinderling/fabscope

